# Multi-Omics Binary Integration via Lasso Ensembles (MOBILE) for identification of context-specific networks and new regulatory mechanisms

**DOI:** 10.1101/2022.07.24.501297

**Authors:** Cemal Erdem, Sean M. Gross, Laura M. Heiser, Marc R. Birtwistle

**Author notes:** Correspondence (L.M.H.); (M.R.B.).

## Abstract

Cell phenotypes are dictated by both extra- and intra-cellular contexts, and robust identification of context-specific network features that control phenotypes remains challenging. Here, we developed a multi-omics data integration strategy called MOBILE (Multi-Omics Binary Integration via Lasso Ensembles) to nominate molecular features associated with specific cellular phenotypes. We applied this method to chromatin accessibility, mRNA, protein, and phospho-protein time course datasets and focus on two illustrative use cases after we show MOBILE could recover known biology. First, MOBILE nominated new mechanisms of interferon-γ (IFNγ) regulated PD-L1 expression, where analyses suggested, and literature supported that IFNγ-controlled PD-L1 expression involves BST2, CLIC2, FAM83D, ACSL5, and HIST2H2AA3 genes. Second, we explored differences between the highly similar transforming growth factor-beta 1 (TGFβ1) and bone morphogenetic protein 2 (BMP2) and showed that differential cell size and clustering properties induced by TGFβ1, but not BMP2, were related to the laminin/collagen pathway activity. Given the ever-growing availability of multi-omics datasets, we envision that MOBILE will be broadly applicable to identify context-specific molecular features associated with cellular phenotypes.

**Graphical Summary:** 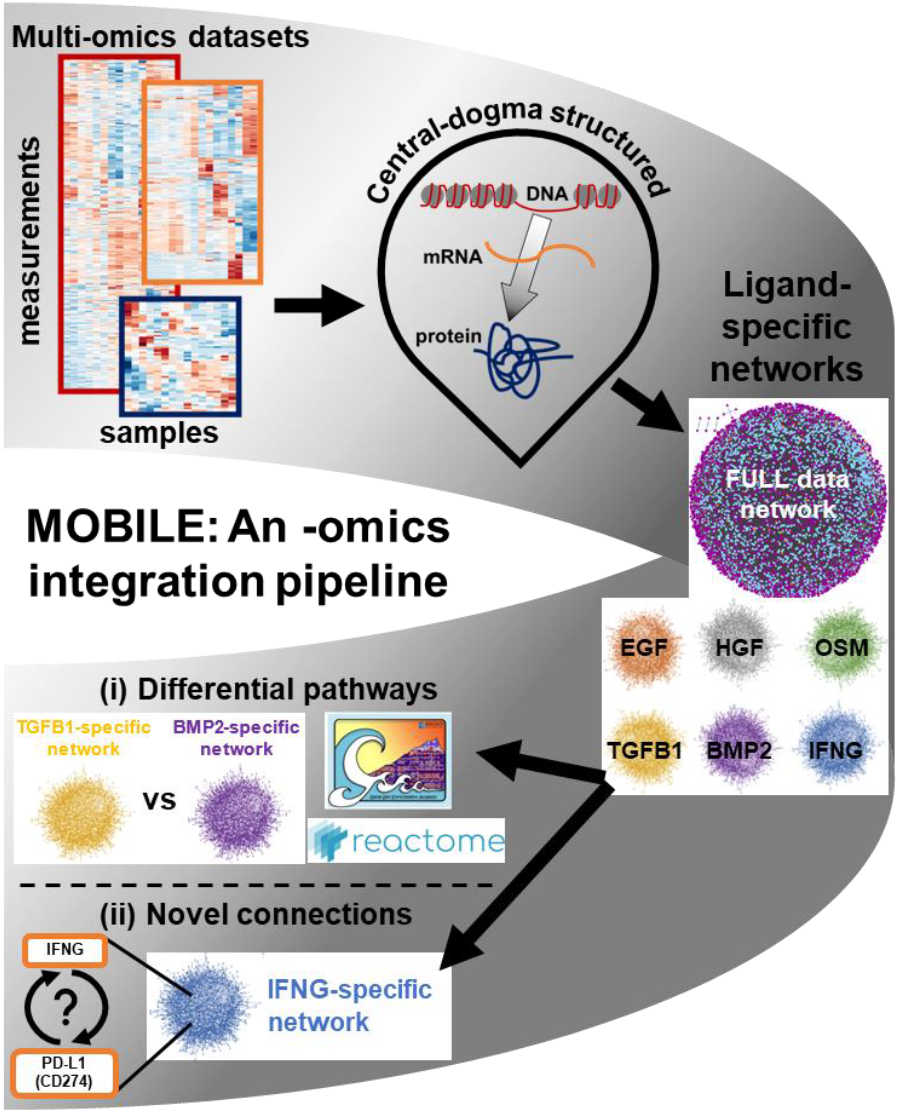

*Multi-Omics Binary Integration via Lasso Ensembles (MOBILE) pipeline yields statistically robust, context-specific association networks:* The MOBILE pipeline integrates omics datasets in a data-driven, biologically-structured manner. The pipeline outputs are gene-level, contextspecific association networks. These association networks nominate differentially enriched pathways, subnetworks, and new connections. Broadly applicable to find condition specific networks using multi-omics datasets.

## Introduction

The availability of large-scale multi-omics datasets across cell types, tissues, and organisms is rapidly increasing with the development of new technologies (1–7). The challenge now is in analyzing them, not individually, but rather together to leverage the fact that signals are encoded across multiple modalities (8–11). Examples of individual methods include: principal components analysis (12), statistical approaches (13–15), clustering (16–19), unsupervised learning (20), supervised learning (21–23), and machine learning (10,24). Often, such analyses are used to generate networks, where genes (or other biomolecules) are nodes, and edges between them denote statistical or functional relationships. Integrating data from multiple modalities can give rise to novel biological insights that cannot be obtained by analyzing single datasets alone (25–27). To date, data integration methods have produced systems-level biological insights including genetic and protein-protein interactions (28–30), disease-gene relationships (24,31), drug response predictions (30,32) and context-specific associations (29,31,33).

A central problem in network biology is understanding what aspects of networks are “context-specific”. Here we define context-specificity as a biological relationship (i.e., edge in a network) that applies only to a certain cell type, stimulatory ligand perturbation, extracellular matrix component, or time point. Data integration may have a significant impact on understanding such context-specific biology, since previous one-at-a-time methods struggle due to the nature of the input data available (8). For instance, why does insulin, but not insulin-like growth factor I (IGF1), regulate glucose metabolism despite their textbook signaling pathways being essentially identical (34–36)? Why does IGF1 activate ERK or AKT signaling only in certain cell types (37,38)? Identifying and studying such context-specific knowledge is important because, first, most diseases have tissue-specificity, and context-specific knowledge could complement available omics data in discovering disease-gene associations (39). Secondly, this enables understanding of how different cell types respond uniquely to varied perturbations, which is key to targeted therapeutic intervention.

There are currently two broad classes of approaches for data integration; one using prior literature / network knowledge as an essential first step (28–33,40–43), and the other not. For example, a prior-knowledge informed network analysis may limit the exploration to curated pathways and remove analytes (measured features) with no known interactions, or may impose biological structure into the underlying network models (29,30). These literature-driven approaches assemble available tissue-specific expression data into gene correlation networks (30,44,45) or overlay the data on global (non-specific) interaction networks (9,28,46). For instance, employing a network/graph theory-based approach coupled with prior literature information, differential network analysis (46–49) tools showed promise in identifying context-specific knowledge (50–59) and helped highlight genes and pathways for clinical impact (11,49,54). While informative, the drawback of such “literature-first” data integration methods is that they only allow study of known connections. This is limiting because a substantial number of regulatory interactions between proteins, mRNAs, micro RNAs, metabolites, and transcription factors are not annotated in the literature (30,46,60). Importantly, these approaches cannot identify novel interactions or associations in the datasets.

The second class of data integration approaches are prior knowledge agnostic, which circumvents the limitations of literature-first methods (24,61–64). These methods rely on extracting features from the input data and constructing models to discriminate between conditions. For instance, Zhang et al. used deep learning to integrate gene expression and copy number variance data from two different databases to identify distinct prognostic subtypes (24). However, the trade-off of these methods is that they are more challenging to interpret because they lack direct incorporation of a biological structure (e.g., central dogma) into the data integration methodology, and also do not take advantage of the wealth of available prior knowledge (11,65,66).

Despite progress, there remains a need for new tools and methods for biologically informed multi-omics integration without literature-driven pre-selection. Here, we introduce **M**ulti-**O**mics **B**inary **I**ntegration via **L**asso **E**nsembles (MOBILE) to integrate multi-omics datasets and identify context-specific interactions and pathways. Our approach does not eliminate data based on prior knowledge and also uses well-established central dogma structure to aid biological interpretation. Robust associations are inferred between pairs of chromatin accessibility regions, mRNA expressions, and protein/phosphoprotein levels. By imposing this high-level structure, MOBILE is neither network structure agnostic (for better interpretability) nor heavily prior-knowledge bound (for higher rates of novelty). We demonstrate the method using a recent multi-omics dataset generated by the NIH LINCS Consortium (67). In that project, we profiled non-tumorigenic breast epithelial MCF10A cells and collected proteomic, transcriptomic, epigenomic, and phenotypic time courses in response to six growth factor perturbations (EGF, HGF, OSM, IFNγ, TGFβ1, and BMP2; synapse.org/LINCS_MCF10A). We apply MOBILE to this dataset and obtain candidate context-specific associations. We then use these associations (i) to propose novel sub-networks of regulation for therapeutically important genes and (ii) to identify pathways preferentially activated by pairs of ligands from similar signaling families. First, MOBILE identifies new regulatory mechanisms for IFNγ-controlled PD-L1 expression that have independent literature support. Secondly, MOBILE reveals and independent experiments validate that TGFβ1 but not BMP2 induces laminin pathway genes (especially laminin 5), causing stronger cell-to-cell and cell-to-surface adhesion through interactions with extracellular collagen, which leads to larger and more separated cells. The biologically structured and data-driven MOBILE pipeline outlined here is widely applicable to integrate omics datasets for extracting context-specific network features and insights.

## Results

### A multi-omics LINCS perturbation dataset for integrative analysis

The NIH LINCS Consortium recently released a unique and comprehensive multi-omics dataset (synapse.org/LINCS_MCF10A). This data set consists of molecular and phenotypic responses of MCF10A cells to multiple ligand perturbations over time (67). Spanning a compendium of canonical receptor signaling classes, EGF, HGF, and OSM induced growth while BMP2, IFNγ, and TGFβ1 inhibited growth. The cellular responses were measured using live-cell imaging, immunofluorescence (IF), and cyclic immunofluorescence (68). The bulk molecular responses were assessed across five platforms. The proteomic assay was reverse phase protein array (RPPA (69)), where specific phospho- or total protein levels were measured at 1, 4, 8, 24, and 48 hours. The RNAseq transcriptomic dataset was single-end sequencing at 24 and 48 hours. Chromatin accessibility was profiled by Assay for Transposase-Accessible Chromatin using sequencing (ATACseq), also at 24 and 48 hours after stimulation. A pretreatment (T0 control) was quantified for all assay types. Additionally, the dataset included global chromatin profiling (GCP) and L1000 transcriptomics readouts (70,71). Overall, the MCF10A dataset provides an excellent template for applying the proposed data integration strategies, namely ATACseq, RNAseq, and RPPA as “big data” to be integrated, and the live-cell imaging / IF as assays informing associated cellular phenotypes.

### The MOBILE Integrator

We here integrated data from this LINCS dataset to identify context-specific pathways and novel regulatory mechanisms that control cellular phenotypes. We reasoned that a data-driven approach whose structure was supported by biological organization could be valuable. The overall approach is summarized in Fig. 1 and presented in greater detail in Fig. 2. The availability of epigenomic, transcriptomic, and proteomic datasets inspired a central-dogmatic view for data integration (Fig. 1). For compatibility across datasets and operability of the method, the MOBILE pipeline input included all three datasets with all ligands at 24 and 48 hours only.

**Fig. 1.**
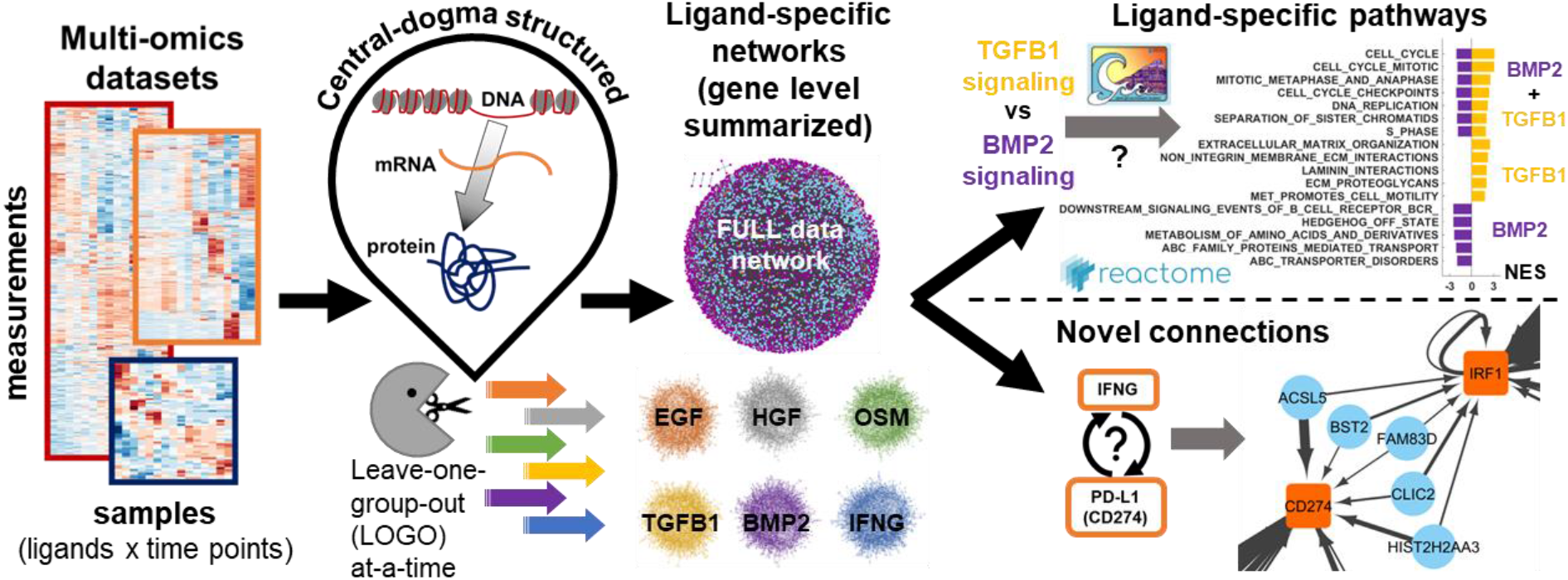
Multi-Omics Binary Integration via Lasso Ensembles (MOBILE) pipeline yields statistically-robust, ligand-specific association networks. The MOBILE data integrator combines multi-omics, multi-assay datasets in a data-driven and central-dogmatic way. By leaving each ligand condition out from the input at-a-time, the pipeline outputs robust ligand-specific association networks. These gene-level networks are used to infer differentially enriched pathways and to find novel regulatory sub-networks.

**Fig. 2.**
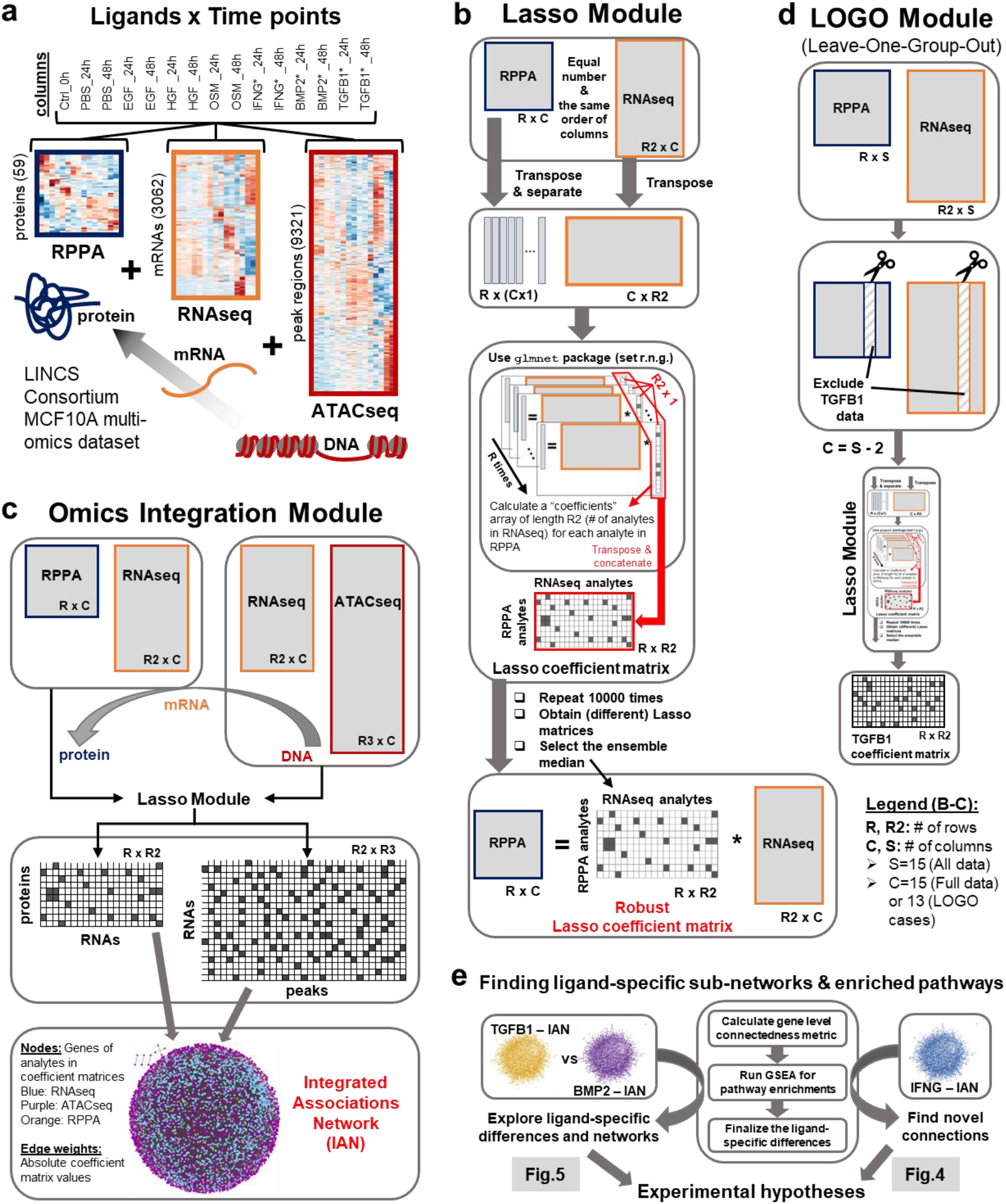
The MOBILE Integrator pipeline transforms input data into gene-level association networks. **a** The NIH LINCS MCF10A omics datasets include proteomic (RPPA), transcriptomic (RNAseq), and epigenomic (ATACseq) assays measured for six ligands + control at 24 and 48 hours. The number of analytes retained after data pre-processing are shown in parentheses in y-axis. The heatmaps shown are actual results from hierarchical clustering of rows. The data show that most ligand conditions at two time points are more similar to each other than to another ligand condition at the same time point. **b** The Lasso module is used to integrate omics datasets one pair at a time. The associations between peaks of open chromatin regions (ATACseq) and mRNA levels (RNAseq) and between mRNA levels and protein levels (RPPA) are calculated separately. The two assay input matrices are structured in a way that yields a robust Lasso coefficient matrix, which contains association coefficients between analytes of the two input matrices. The pipeline iteratively applies the lasso algorithm to each row of the input data matrix on the left-hand side and calculates one row of the Lasso coefficients matrix. Minimizing the mean squared error and penalizing the L1 norm, only a few non-zero coefficients are selected at each iteration. The resulting coefficient vectors for every analyte are concatenated to obtain a single Lasso coefficient matrix instance. Each non-zero element of this matrix represents the regression coefficient obtained for the effect of the analyte of right-hand-side input matrix on the level of analyte in the left-hand-side input matrix: the effect of mRNA levels on (phospho)protein levels or the effect of open chromatin peak regions on mRNA levels. Ten thousand instances of the Lasso coefficient matrix are generated, starting each with a different seed for random number generator. The coefficients that appear in at least half the matrices (>5000 times) are considered robust and the representative matrix with the largest number of robust coefficients is selected as the robust Lasso coefficient matrix. The coefficients (non-zero elements) in this matrix are called associations for the remainder of this work. **c** The robust Lasso coefficient matrices of two different pairs of datasets (i.e. RPPA+RNAseq and RNAseq+ATACseq) are combined to generate Integrated Association Networks (IANs) for each ligand condition. These gene-level networks represent robust, statistical associations inferred from multi-omics datasets, offering a new hypotheses generation tool to look for ligand or gene-set specific sub-networks. Node colors represent; blue:RNAseq, purple:ATACseq, and orange:RPPA and the edge widths correlate with the magnitude of the association coefficients. **d** We employ a leave-one-group-out (LOGO) analysis to find associations that show-up or disappear depending on exclusion/inclusion of that specific condition. We exclude one set of ligand conditions (24 and 48 hours) from the input matrices, run the Lasso module (**b**) with a smaller number of columns, and obtain a new robust Lasso coefficient matrix. Such matrices are called ligand-specific coefficient matrices. Comparing the resulting matrix from the LOGO module to the FULL-data Lasso coefficient matrix, we determine coefficients dependent on the existence of the corresponding ligand data. Finally, we combine the “ligand”-dependent coefficients with the coefficients that disappear from the FULL-data matrix (when “ligand” conditions are excluded from training) to create the final ligand-specific associations list. **e** Using the corresponding IANs, we generate a novel interactions sub-network of IFNγ and PD-L1 relationship and explore TGFβ1 and BMP2 enriched pathways.

Following the central dogma that information flows from DNA to RNA to protein, we paired ATACseq-RNAseq and RNAseq-RPPA matrices. First, we calculated robust and parsimonious statistical associations between features of input data (Fig. 2a, see Methods) using replicated penalized regression models (Fig. 2b, Supplementary Data 1 and 2). We applied Lasso (least absolute shrinkage and selection operator (37,72)) regression to infer sets of sparse matrices that can be interpreted as statistical networks connecting biochemical species, such as mRNAs, chromatin peaks, or total protein and phosphorylation levels. The repetitive application of Lasso, called the Lasso module, yielded an ensemble of such matrices, from which we picked the matrix with the greatest number of robustly inferred associations as the robust associations matrix. When we used all the ligand conditions from the LINCS dataset as input, the Lasso module output is called the FULL-data matrix. To finalize the data integration and generate data-driven networks, we merged the robust associations matrices obtained from RPPA+RNAseq and RNAseq+ATACseq input pairs into an Integrated Association Network (IAN) (Fig. 2c). The IANs were coalesced gene-level networks, where nodes represent genes of the assay analytes (genes from input matrix rows), and edges represent robust Lasso coefficients calculated between the analyte levels (Supplementary Fig. 1).

We then systematically excluded different ligand conditions (both 24- and 48-hr data) from the training input and ran the LOGO (leave-one-group-out) module (Fig. 2d). We hypothesized that the robust associations that change as a result of this LOGO analysis will have information regarding the context-specificity of the left-out ligand condition. The ligand-specific IANs together with the FULL IAN were the major data-integration products of the MOBILE. Comparison of pairs of ligand IANs nominated differentially activated pathways and novel, ligand-specific regulatory mechanisms spanning DNA states to protein levels. To extract this information, we performed gene-set enrichment analysis (GSEA (73)), which revealed significantly enriched pathways based on the IAN gene lists. Here, using the Reactome database (74) enabled us to identify pathways linked to ligand-specific IANs. By analyzing the enriched pathways and relating them to the corresponding phenotypic responses, we nominated ligand-specific pathways and novel edges playing a role in the distinct phenotypes (Fig. 2e).

### MOBILE identifies known biology

We investigated the robustness of the MOBILE predictions by performing a gene-set enrichment analysis (GSEA) for pathways using the FULL and ligand-specific integrated associations networks (IANs) and asking whether our approach can capture canonical biological observations. First, we identified ligand-dependent association lists by comparing each ligand IAN to the FULL IAN. Next, these association lists are coalesced into gene-level networks and the nodes are ranked based on the sum of edge weights (association magnitudes) entering that node (Supplementary Data 3). By running the GSEA on these eight (FULL, PBS, EGF, HGF, OSM, IFNG*, BMP2*, TGFB1*) pre-ranked gene-lists, we saw that the top two/three enrichments (p<0.05 and FDR<0.1) are cell cycle pathways and are significantly enriched for all conditions (Fig. 3a, Supplementary Data 4). Indeed, eight of the top 15 pathway enrichments across conditions are cell cycle related, affirming the fact that the LINCS dataset was generated using combinations of pro/anti-growth factors and cells continue to grow after all perturbations (Supplementary Fig. 2 and Supplementary Data 5).

**Fig. 3.**
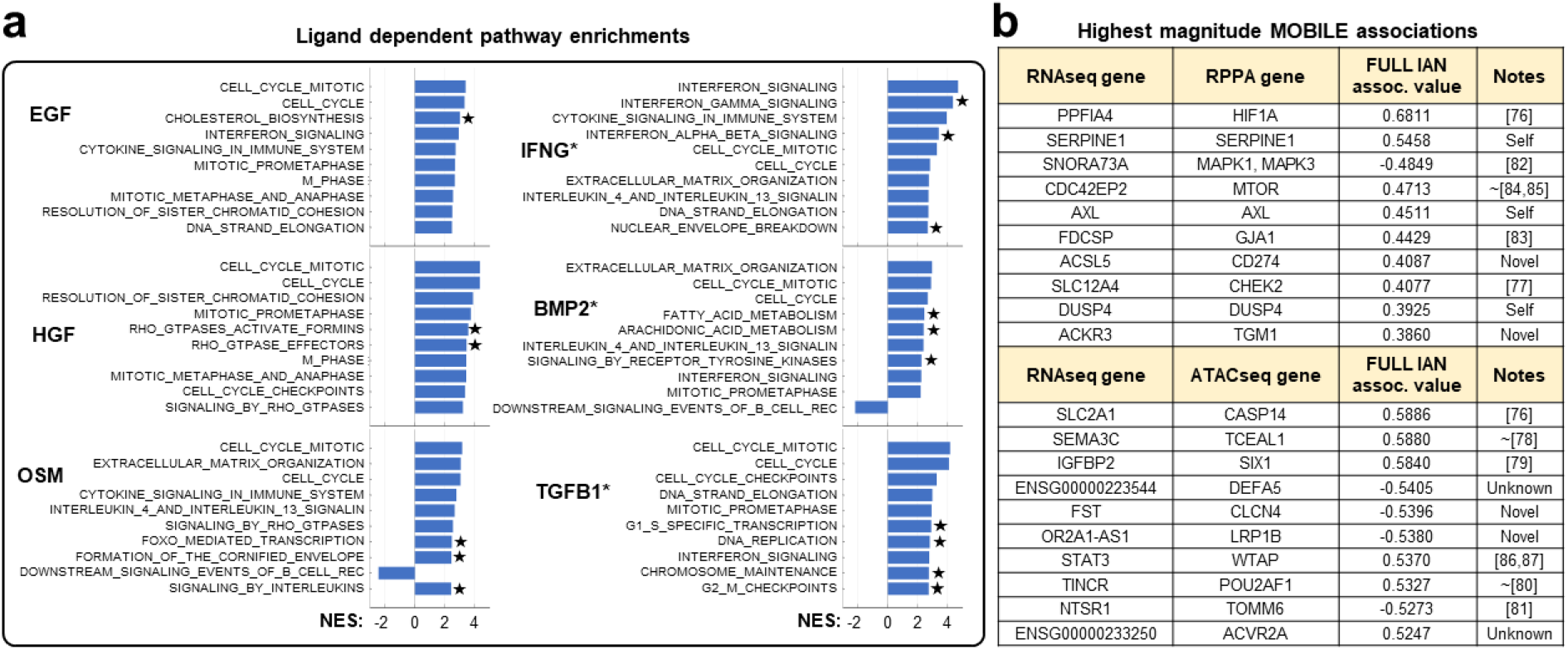
MOBILE inferred IANs are enriched for canonical pathways and the top associations are literature verified interactions. **a** Gene set enrichment analysis using genes of ligand dependent associations reveal cell cycle, interferon, and cytokine signaling pathways. Only top 10 significantly enriched pathways are shown for each ligand perturbation (p-values < 0.05, FDR<0.1). The bar height corresponds to the normalized enrichment scores (NES). Stars (⭐) denote pathways enriched specifically for the corresponding ligand condition when only considered top 10 pathways shown. * denotes conditions with additional EGF treatment. **b** MOBILE finds known and novel associations between proteins, transcripts, and chromosomal region genes. Top 10 magnitude-wise associations between RPPA-RNAseq and RNAseq-ATACseq of the FULL IAN are presented. More than half of the associations have prior literature evidence, while some are Self associations of the same gene in different assays. Two associations include non-annotated gene products and labeled as Unknown. At least four of the associations are Novel predictions of the MOBILE pipeline. ~ denotes references not showing direct (causative) relationship between the genes but co-mentioning them as biomarkers of different cancer subtypes or with relationships of candidates’ isoforms. The MOBILE inferred association values are between −1 and 1.

Other highly enriched pathways in the HGF dependent gene-list were Rho GTPase related (Fig. 3a). It was shown before that Rho GTPase activity is required for HGF-induced cell scattering (75). OSM-dependent pathway enrichments included cytokine/interleukin and ECM pathways (Fig. 3a). The top four highly enriched pathways of IFNG* condition were interferon signaling, in line with the fact that IFNγ had a strong signal in the LINCS dataset (67). BMP2-dependent top pathways were ECM, interferon, and interleukin related in addition to the cell cycle. Finally, TGFB1* condition had transcriptional and DNA regulatory pathways enriched (Fig. 3a). These observations confirmed that our approach can recover known biology and also that it can extract meaningful ligand-specific associations.

Next, we asked whether MOBILE-inferred associations (edges) are consistent with prior knowledge. The highest magnitude associations of the FULL analysis are the most robust ones across all perturbations and timepoints (Fig. 3b, Supplementary Data 1 and 2). Among them, the top candidate interaction is the connection between PPFIA4 (Protein Tyrosine Phosphatase Receptor Type F Polypeptide-Interacting Protein Alpha-4) and HIF1A (Hypoxia Inducible Factor 1 Subunit Alpha). The PPFIA4 gene was shown to be upregulated in response to hypoxia (through HIF1) in all types breast cancer cell lines and normal-like epithelial cells, including MCF10A (76). The highest association between ATACseq and RNAseq data was the SLC2A1 (Solute Carrier Family 2 Member 1) and CASP14 (Caspase 14). Interestingly, these two genes were also part of the hypoxia-induced genes list (76). There exists literature evidence for the other highest-ranking associations (Fig. 3b). Some were (i) shown to be part of prognostic markers (SLC12A4-CHEK2 (77)), (ii) differentially expressed together in response to perturbations (SEMA3C-TCEAL1 (78), IGFBP2-SIX1 (79), TINCR-POU2AF1 (80), NTSR1-TOMM6 (81)), and (iii) part of gene signatures for different classes of tumors (SNORA73A-MAPK (82), FDCSP-GJA1 (83)). A few of the associations had related mechanistic interactions as well (CDC42-MTOR (84,85), IGFBP2-SIX1 (79), WTAP-STAT3 (86,87), TINCR-POU2AF1 (80)). The rest of the associations are either Self: same gene, different data type, Unknown: non-curated gene(s), or Novel: no known interactions. These pieces of information from the literature in-part verifies that the MOBILE inferred associations have biological meaning.

### Identification of novel associations between IFNγ stimulation and PD-L1 regulation

After establishing that MOBILE can recapitulate known biological interactions, we asked whether it could identify novel regulatory mechanisms within a single IAN. We focused on IFNγ, which had a strong signal in the LINCS dataset (67) and is a critical part of the immune response within the tumor microenvironment (88,89). The cytokines within the environment, especially IFNγ, can induce transient PD-L1 (gene name: CD274) expression (Fig. 4a) (90–93). PD-L1 is a transmembrane protein that binds to its receptor PD-1 expressed in T-cells and inhibits immunological tumor clearance. Both PD-L1 and PD-1 belong to a class of so-called “checkpoint” proteins (91,94); immune checkpoint inhibitors are a new class of immunotherapeutic anti-cancer drugs (93,95). However, not all cancers or patients are responsive or have drug resistance to such therapy because of high PD-L1 expression variability (depending on tumor stage, tumor site, tumor type). Consequently, predicting tumor responses to PD-1/PD-L1 blockade remains a challenge, and better biomarkers are needed to stratify patients. Therefore, an in-depth understanding of the regulatory mechanism of PD-L1 expression is still needed to provide new immunotherapeutic insights and potentially identify new drugs (96). To investigate this question, we decided to explore novel sub-networks between IFNγ signaling and PD-L1 expression within the data-driven IFNγ integrated associations network.

**Fig. 4.**
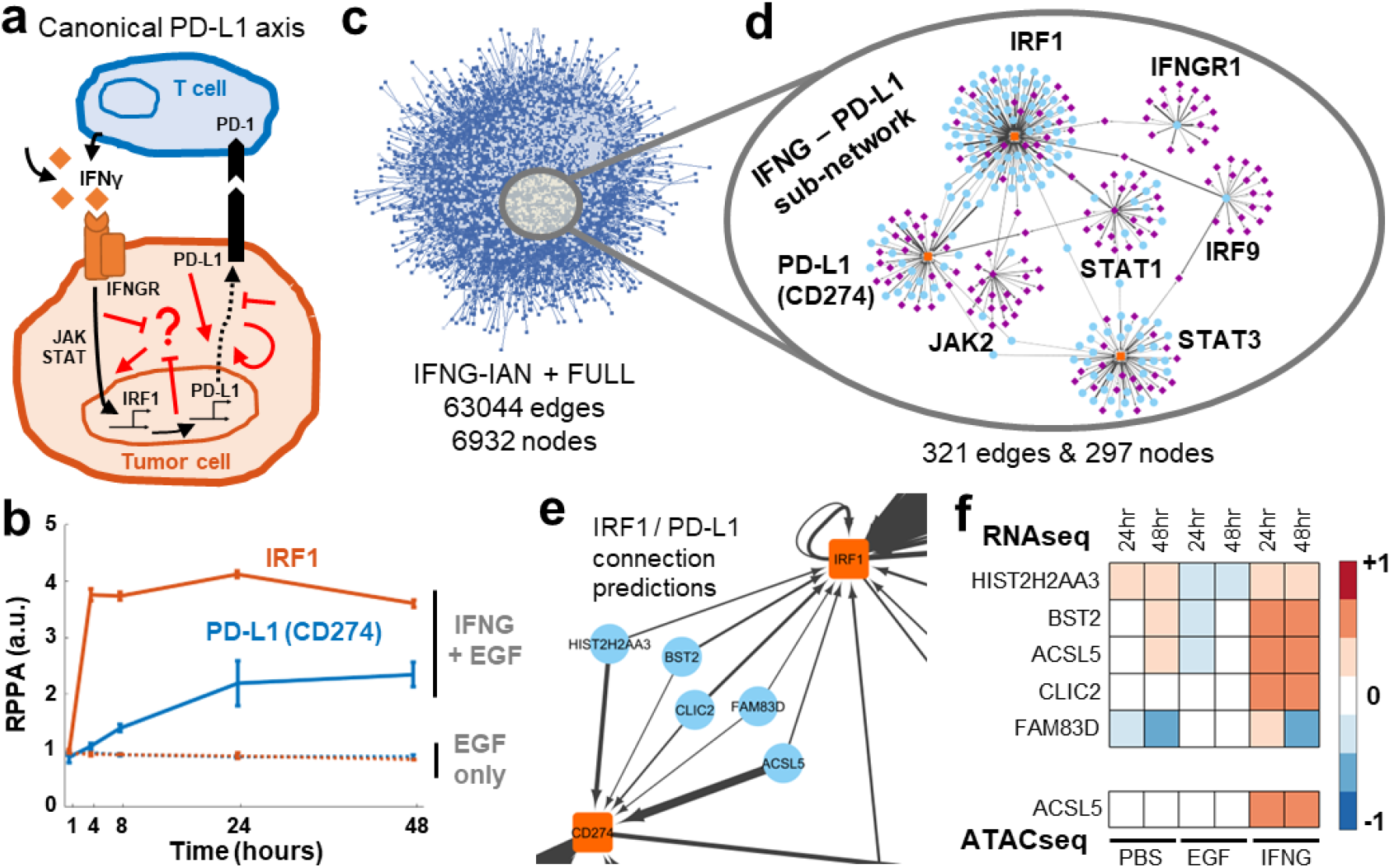
Exploration of single integrated association network reveals new links between IFNγ signaling and PD-L1. **a** IFNγ secreted by T cells induces PD-L1 expression through JAK/STAT/IRF1 and other canonical pathways (black arrows). The PD-L1 on the cell surface then interacts with PD-1 on the immune cells to induce tumor cell death. However, the PD-1/PD-L1 therapy yields inter- and intratumor heterogeneous response and there is a need to identify new non-canonical targets (represented by red arrows). **b** The IFNγ induces IRF1 and PD-L1 production in MCF10A cells. **c** The IFNG associations network (IFNG-IAN) is a data-driven large-scale network of novel connections. The associations are coalesced into gene-level nodes, and associations with greater than 0.01 absolute value are shown. **d** The sub-network of IFNγ – PD-L1 relationship is significantly smaller than the IFNG-IAN. The sub-network is generated by the filtering for 14 genes and has seven hubs (PD-L1 (CD274), IFNGR1, IRF1, IRF9, JAK2, STAT1, STAT3) connected to 290 other genes. Node colors represent; blue:RNAseq, purple:ATACseq, and orange:RPPA and the edge widths correlate with the magnitude of the association coefficients. **e** A closer look at the connections between IRF1 and PD-L1 (CD274) genes shows a breadth of different functional genes. **f** Compared to no or EGF-only stimulation, the connecter genes are upregulated (RNAseq) in IFNγ stimulated condition. Additionally, the ACSL5 gene peak is more accessible similarly to the canonical IFNγ downstream genes (ATACseq). All data shown are from the pre-processed input.

In the LINCS dataset, IFNγ was tested in combination with EGF, so here we compared the IFNγ condition (IFNγ+ EGF) to EGF-only samples. It is known that IFNγ upregulates PD-L1 on cells (97). First, looking at the non-tumorigenic MCF10A cell data, we also verified that IFNγ stimulation upregulated IRF1, interferon regulatory factor, and PD-L1 (CD274 gene) levels, with no expression in the EGF only case in the LINCS RPPA data (Fig. 4b). Next, we identified a set of nine genes (IFNG, IFNGR1, IFNGR2, STAT1, STAT3, JAK1, JAK2, IRF1, IRF9) from the canonical IFNγ pathway (REACTOME R-HSA-877300) and filtered the connections of the genes together with PD-L1 (CD274) and PD-1 (CD279) from IFNG-IAN (Fig. 4c). The resulting sub-network had 297 nodes and 321 edges (Fig. 4d, Supplementary Data 6 and 7). The hubs in that sub-network were from the input gene list, including IRF1, STAT1, STAT3, and PD-L1. We examined the connections between IRF1 and PD-L1 (Fig. 4e) and identified a five-gene set of connectors: BST2, CLIC2, FAM83D, ACSL5, and HIST2H2AA3. The mRNA levels of these five genes were elevated in the IFNγ+EGF condition compared to PBS control and EGF-only samples (Fig. 4f). Of the five genes, we could not find any literature data for two connections (ACSL5 and HIST2H2AA3) and their relationships to IFNγ and PD-L1 function. Therefore, studying these novel connection predictions may reveal further diagnostic and therapeutic targets for immunotherapy. Notably, the remaining three genes had strong literature data. BST2 was recently shown to be part of a gene signature for anti-CTLA4 response in melanoma (98). CLIC2 is co-expressed with PD-L1/PD-1 in breast cancer and is a biomarker candidate for favorable prognosis (99). And although not directly linked to IFNγ/PD-L1 axis, FAM83D was shown to regulate cell growth and proliferation and was implicated as a prognostic marker in breast and gastric cancers (100–102). Moreover, its FAM83A isoform was shown to affect PD-L1 expression (103). Overall, the concordance of these findings with recent literature reinforces the notion that MOBILE-based nomination of novel interactions has biological value.

### TGF-β superfamily members TGFβ1 and BMP2 induce different morphological phenotypes via collagen-laminin signaling

Both BMP2 and TGFβ1 are members of the TGF-β superfamily of ligands and share most downstream pathways including canonical SMAD signaling (104,105). Both ligands induce cell differentiation and show anti-growth/anti-proliferative effects, where SMAD signaling shows immense versatility and specificity, mostly affected by the cross-talk mechanisms and the cellular context (104–112).

Imaging data of cells grown on collagen-coated culture plates from LINCS (67) indicated that BMP2 induces a significantly higher number of cells in clusters compared to that induced by TGFβ1 (Fig. 5a, top). Correspondingly, TGFβ1 induced morphologically larger cells (113,114) that occupy more surface area (Fig. 5a, bottom). So, we used the ligand-specific IANs (Supplementary Fig. 3) and subsequent pathway enrichment analyses to find non-canonical mechanisms that underly the differential phenotypes caused by these two highly similar ligands. We ranked the nodes of the TGFβ1 and BMP2 IANs (Supplementary Data 8 and 9) based on the sum of edge weights entering that node (Supplementary Data 10 and 11) and ran GSEA on these pre-ranked gene-lists (73,115). We then looked for curated Reactome pathways (74) significantly enriched (p<0.05 and FDR<0.1) in either gene-list (TGFβ1 or BMP2 IAN genes) (Fig. 5b). Seven of the enriched pathways under BMP2 and TGFβ1 treatment conditions are shared (Fig. 5c gray circles, and Supplementary Data 12 and 13). The shared pathways are all cell cycle and proliferation related. Multiple pathways are specific to a single condition (Fig. 5b, BMP2: purple and TGFβ1: gold). BMP2 enriched pathways include DNA regulation and G1/S transition. The TGFβ1-only group had DNA damage and ECM regulation-related clusters of pathways. The laminin interactions (REACTOME R-HSA-3000157) pathway was significant among those that were ECM-related. Laminins are a family of proteins that regulate cell-to-cell and cell-to-environment interactions (116–121). It is known that laminins bind to collagen (122), integrin receptors (123), and depletion of ECM laminin or collagen disrupts the cellular attachment (116,124). Differential regulation of such a pathway might explain the phenotypic differences of TGFβ1 and BMP2 phenotypes and offer a candidate mechanism to explore further for enlarged (more spreading and stretching) cells specific to the TGFβ1 response. Specifically, that only TGFβ1 induces laminin gene expression that then interacts with the collagen-coating of the culture plate, which induces tighter cell-to-cell and cell-to-ECM interactions.

**Fig. 5.**
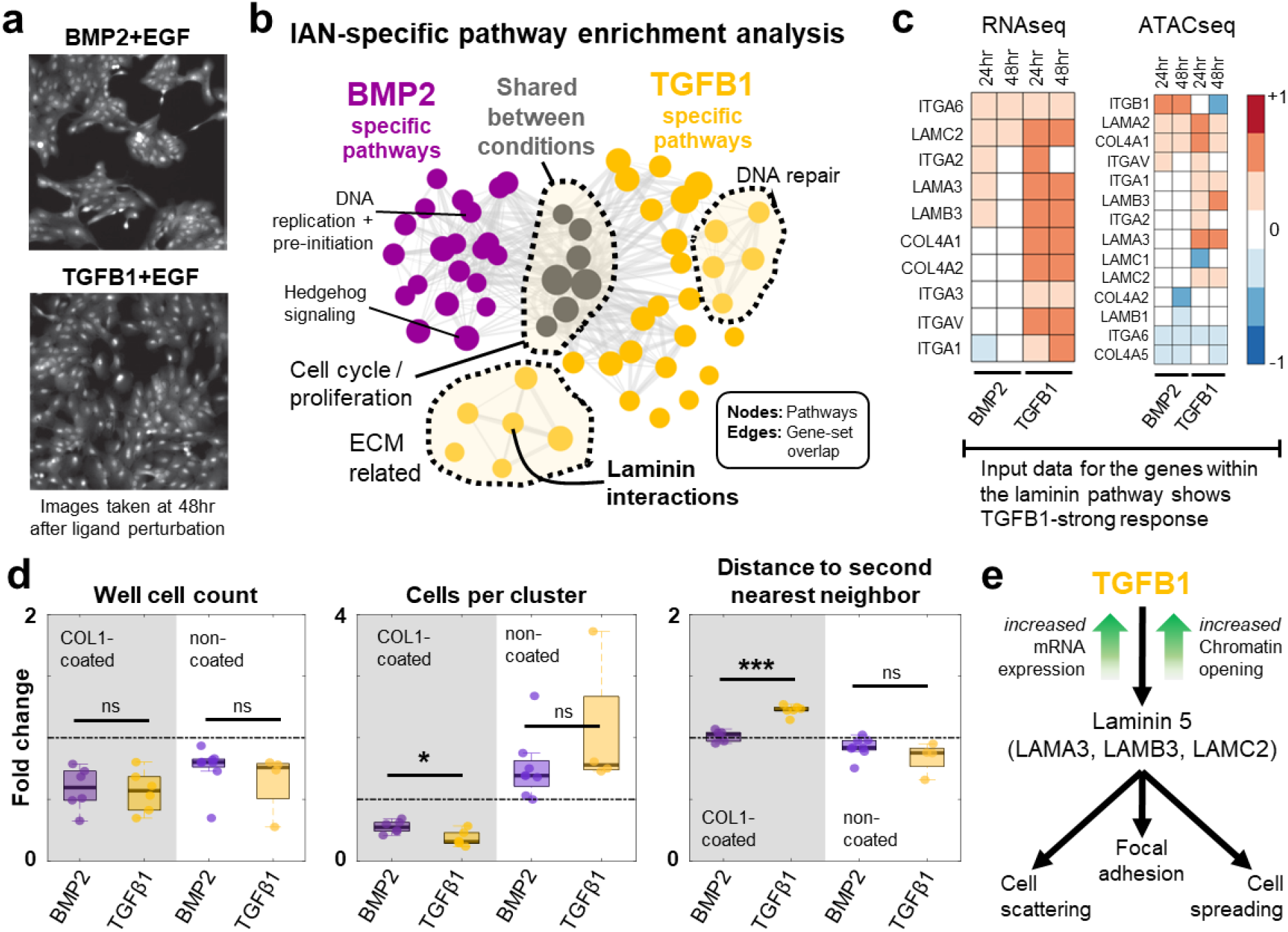
TGFβ1 and BMP2 induce different morphological phenotypes by differentially activating collagen-laminin signaling. **a** TGFβ1 and BMP2 induce phenotypically different responses as seen in the immunofluorescence images 48 hour after ligand treatment (adapted from (67)) **b** The ranked gene lists of the ligand-specific (TGFβ1 or BMP2) networks are used to find enriched lists of curated Reactome pathways. Both conditions are enriched in cell cycle and proliferation related pathways (gray circles). TGFβ1 alone is shown to regulate ECM related pathways whereas BMP2 network genes are linked to other membrane receptor related signaling pathways and cell cycle checkpoints. Node size is proportional to the enrichment scores and edge widths represent the number of overlapping genes within the connected pathways. **c** The laminin pathway regulates cell-to-cell and ECM adhesion of cells and is shown to be upregulated by TGFβ1 alone. The expression levels of the genes (left) within the pathway are upregulated by TGFβ1. The gene regions (peaks, right panel) of the laminin genes are more accessible in TGFβ1 condition. Laminin 5 is the major secreted complex consisting of LAMA3, LAMB2, and LAMC2. Transcriptomic and epigenomic data shown are from the pre-processed input. **d** Quantification of the images in (**a**) shows that both BMP2 (purple) and TGFβ1 (gold) induce similar anti-growth responses but yield significantly different microenvironmental and spatial characteristics. The metrics shown are normalized to the mean of EGF-only treatments at 48 hours (black dashed line), where both ligands inhibit cell proliferation and cause less number of cells (cell counts, left box plot, shaded region). However, BMP2 induced larger number of cells in clusters compared to TGFβ1 (middle box plot, shaded region, p-value = 0.0175). TGFβ1, on the other hand, caused cells spread and have longer distance to their second nearest neighbors (right box plot, shaded region, p-value = 9.4032E-06). To confirm MOBILE predicted effect of collagen-laminin interactions for such differences in ligand responses, we performed validation experiments where cells are grown in non-collagen-coated, regular tissue-culture treated plates. The results show phenotype reversal of cells-per-cluster and distance-to-second-neighbor (unshaded regions, middle and right box plots), with non-significant differences between BMP2 and TGFβ1. The box edges correspond to 25^th^-75^th^ percentiles, the horizontal black lines represent the median, and the dots are individual data points. The whiskers extend to the extreme (non-outlier) data points. ns: not significant (p-value > 0.05). * represents p-value < 0.05, *** p-value < 0.001. Significance tested using Student’s t-test with unequal variance and both tails. **e** The schematic of TGFβ1 specific regulation of a non-SMAD pathway, inferred by the MOBILE Integrator pipeline. TGFβ1 induces laminin pathway gene expression and leads to cell scattering and cell spreading with larger cells, when compared to BMP2 stimulation.

Next, to initially confirm the activity of the laminin pathway under TGFβ1 but not BMP2 stimulation, we referred back to the mRNA levels (the MOBILE input data) of the laminin interactions pathway genes (Fig. 5c) and saw TGFβ1-upregulated transcription compared to BMP2. The mRNA levels of laminin, collagen, and integrin subunits were elevated at 24 and 48 hours (Fig. 5c, left), and corresponding transcription binding sites were more accessible in TGFβ1 stimulated cells (Fig. 5c, ATACseq data, right).

In the original dataset, cells were cultured and treated on collagen-coated plates (67) and both ligands inhibited cell proliferation compared to EGF-only control (Fig. 5d, left panel). Additionally, BMP2 induced more cells in clusters compared to TGFβ1 (Fig. 5d, middle panel) but TGFβ1 was shown to cause significantly larger distances to second nearest neighbors (Fig. 5d, right panel). To test the hypothesis that TGFβ1-specific phenotypes depend on activation of laminin-collagen (in ECM) interactions (Fig. 5e), we cultured cells on non-collagen-coated plates and stimulated them with TGFβ1 or BMP2 (+EGF, as done in the LINCS dataset (67)). Removal of the collagen caused a reversal of TGFβ1-induced phenotype. We saw that the number of cells per cluster as well as cell centroid distances to second nearest neighbors are similar in BMP2 and TGFβ1 stimulated cells in the absence of collagen-coating (Fig. 5d, middle and right panels). Overall, these analyses confirmed that MOBILE identified a context-specific network that explains differential phenotype between two highly similar ligands.

Finally, although MOBILE analysis identified the laminin pathway as potentially explanatory for differential phenotype, we wondered whether such a conclusion could be reached by standard differential expression analysis. Considering the same list of pre-filtered 3062 transcripts that were input to the MOBILE pipeline, we determined BMP2 or TGFβ1 up and down regulated genes at 24- or 48-hour conditions. Next, we ranked the genes based on the fold-change (BMP2 vs TGFβ1 or TGFβ1 vs BMP2) and, similar to post-MOBILE enrichment analysis, we ran GSEA on these pre-ranked lists of genes (Supplementary Data 14). We then compared MOBILE results with the differential expression analysis results. The BMP2 up-regulated, TGFβ1 up-regulated, BMP2 down-regulated, and TGFβ1 down-regulated gene lists alone did not yield any significant pathway enrichments (p<0.05 and FDR<0.1). However, we obtained a single significantly enriched pathway (REACTOME Extracellular Matrix Organization, R-HSA-1474244) when we looked at the combined list of TGFβ1 up- and down-regulated genes. The combined BMP2 regulated gene list yielded 10 significantly enriched pathways including ECM Organization (Supplementary Data 15), whereas the MOBILE pipeline yielded 39 (TGFβ1, Supplementary Data 12) and 20 (BMP2, Supplementary Data 13) enriched pathways using TGFβ1- and BMP2-specific IANs. In summary, we conclude that the MOBILE inferred ligand-specific association networks and their analyses extract more information on the differential pathway enrichments compared to standard methods.

## Discussion

Moving away from the single modality analysis of “multi-omics” datasets, data integration methods are becoming more widely available due to their ability to extract multi-scale information (i.e., genomic to proteomic). Some data integration methods are exclusively based on prior-pathway knowledge, while others utilize such knowledge to infer human-interpretable associations. Here we introduced the MOBILE pipeline to integrate and analyze multi-omics datasets in a data-driven way. With a central-dogmatic hierarchy, the method finds statistically robust integrated association networks (IANs) between epigenomic, transcriptomic, and proteomic analytes that is ultimately human interpretable. We explored ligand-specific IANs obtained via leave-one-group-out (LOGO) analysis– for non-canonical connections with novel immunotherapeutic potential and differentially activated pathways to discriminate between highly similar ligand-receptor responses.

The MOBILE pipeline can prioritize context-specific, differentially activated pathways and mechanisms. The pipeline finds statistical associations where the LOGO analysis is the part that provides context-specificity. By holding out data from each ligand one at a time (24- and 48-hour time points), associations that depend on the held-out data are inferred and cataloged as ligand-dependent. These ligand-dependent associations are the core of MOBILE integrator, enabling exploration of single ligand-specific and differentially enriched pathways between multiple ligand conditions. We expect that the exact method used for forming statistical associations (replicated LASSO) is not particularly essential, and other methods may be substituted so long as it is robust and can make use of the information in the datasets (10,24,28,30,44–46). However, by imposing matching time points, we miss time-lagged associations between mRNAs and proteins due to their temporal ordering. Thus, a next step for the MOBILE pipeline could be to become more flexible with respect to different time points for different assays and fully exploit temporal dependence information for inference of associations.

The above-mentioned matching column order for input matrices requirement of the MOBILE pipeline in this work does not restrict users to study only time points x ligand conditions. For instance, Erdem et al. showed that paired input data matrix pairs at different time points (rows: proteomic measurements, columns: different cell lines) could infer robust, time-dependent associations between proteins and phospho-proteins (37). Another way to utilize the MOBILE pipeline is to find cell line-specific association networks by considering datasets like Cancer Cell Line Encyclopedia (CCLE), where hundreds of cell lines were characterized with molecular and functional assays (2,125). MOBILE could be set up where columns represent different cell lines, and the matrix pair are proteomic and transcriptomic data. By imposing different higher-level hierarchies for the MOBILE pipeline, researchers can explore different types of context-specificity by using data from either single or multiple assays.

Nevertheless, the lists of associations generated by MOBILE are all data-driven experimental candidates to study ligand-specific linkages between genes and gene products. Especially, the highest magnitude associations could be starting points for new experiments to explore crosstalk mechanisms or unknown links in the literature. For instance, by analyzing the IFNγ-specific network only, we hypothesized new regulatory mechanisms of PD-L1, a critical immunotherapeutic target. Of the five MOBILE-hypothesized connector genes (BST2, CLIC2, FAM83D, ACSL5, and HIST2H2AA3) between IRF1 and PD-L1, BST2 was recently recognized as part of an immune/tumor related signature that is significantly associated with the overall survival of skin cancer patients (98). Specifically, the BST2 gene signature predicted response to a CTLA4 antibody called ipilimumab, suggesting a mechanistic involvement in tumor progression. Secondly, CLIC2 was shown to be co-expressed with PD-L1 and PD-1 and act as a good prognosis marker with higher rates of tumor-infiltrating CD8+ T cells in breast cancer patients (99). Finally, FAM83D was shown to be a potential oncogene with high expression levels associated with poor breast cancer prognosis (100). Moreover, FAM83A (an isoform of FAM83D) was shown to drive PD-L1 expression and be correlated with poor lung cancer prognosis (103). These results suggest that context-specific gene-gene associations identified through MOBILE are potential biomarkers for prognosis and patient response to immune checkpoint inhibition.

Another key capability of MOBILE pipeline is the IAN-comparative analysis. For instance, we hypothesized and then experimentally validated that the phenotypic differences between TGFβ1 and BMP2 perturbations were caused by cell-ECM interactions, specifically laminin-collagen. When we replicated LINCS experimental conditions in tissue culture plates without collagen coating, the two ligands induced similar responses (Fig. 5d), where both ligands were previously known to induce cell differentiation, inhibit cell proliferation, and signal through similar canonical pathways (105–107). Additionally, it was shown before that laminin/collagen pathway inhibition leads to cell-ECM attachment disruption (116,124). However, there are other TGFβ1-specific ECM-related pathways (Fig. 5b) and genes that could be further explored for differences between BMP2 and TGFβ1 conditions. Similarly, studying other ligand-IAN pairs (e.g., EGF vs HGF, IFNγ vs TGFβ1, OSM vs IFNγ) could suggest additional data-driven hypotheses.

The MOBILE pipeline here infers robust associations between genes and genes products without prior network knowledge input, enables generation of context-specific, gene-level networks of different biological modalities in a data-driven way, and provides exploration of these networks in a single or paired fashion to pinpoint differentially activated pathways. We believe the freely-available MOBILE pipeline will be broadly helpful in extracting novel context-specific insights from multi-omics datasets to help answer targeted biological questions.

## Methods

### Computational methods

#### The Lasso module

The multi-omics datasets from the LINCS consortium (67) are pre-processed using a raw variance filter to retain only 10% (RNAseq, ATACseq) and 20% (RPPA) highly variant analyte measurements across median summarized Level 4 data (Fig. 2a, synapse.org/LINCS_MCF10A, Source Data). The proteomic (RPPA), transcriptomic (RNAseq), and epigenetic (ATACseq) datasets are integrated with a central dogmatic view, such that pairs of RPPA+RNAseq and RNAseq+ATACseq data matrices are run through the Lasso module (Fig. 2b-d). The steps of the algorithm (37) are:

1. We use glmnet package for lasso regression (126). The module takes two input matrices, Y=left hand side and X=right hand side, and our goal is to calculate matrix β in *Y* = *β* · *X* + *δ*. The number and the ordering of columns in input matrices should be equal (Fig. 2b). The rows are assay analytes measured and columns represent ligand/timepoint conditions.
2. The matrices are column centered, row-centered, and row normalized. The preprocessing makes sure that the algorithm is not biased towards high magnitude analyte measurements but focused on analyzing based on the shape of measurements across conditions. It also sustains that the offset value is moved towards zero.
3. We set cross-validation parameter of `glmnet` package to 4 and turned-off the input data standardization option.
4. Next, both matrices are transposed and the transposed Y matrix (Y’) is separated into column vectors.
5. For each column k of the Y’ (or each row of input matrix Y), a set of lasso regression coefficients are calculated using `glmnet` package. With every iteration, we obtain one row of the final coefficient matrix β and an offset value δ, which is negligible in this case (values less than 10^-7^). We minimize the quantity:

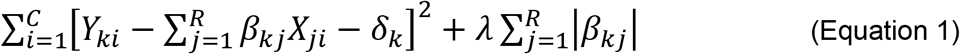
6. The λ factor (Eqn. 1) is estimated via the inherent cross-validation step of the glmnet package. In short, a set of different λ values are tested resulting in different sets of lasso coefficients, each with potentially different number of non-zero coefficients.
7. We select the set of lasso coefficients (i.e., the lasso coefficient vector) corresponding to the minimum estimation error.
8. Repeating steps 5-7 as many times as the number of input Y rows, we obtain R-many Lasso coefficient vectors, each R2-long.
9. We concatenate the R-many R2-long vectors to obtain a Lasso coefficient matrix.
10. We repeat the (3–9) steps 10000 times to obtain an ensemble of Lasso coefficient matrices. We start each estimation with a different seed for random number generator.
11. We then find coefficient indices that appear at least half of the time (5000 times) and select the matrix with highest number of such coefficients. This matrix is called the Robust Lasso coefficient matrix and used for the rest of the analyses.
12. The final matrix (β) sustains the equality *Y* = *β* · *X* and contains association weights relating the analyte levels of the two input matrices.
13. When the input data contains all experimental conditions, we named the resulting robust Lasso coefficient matrix as the FULL-data matrix.
14. We do steps 1-13 for (i) RPPA (matrix Y)-RNAseq (matrix X) and (ii) RNAseq (matrix Y)-ATACseq (matrix X) input data matrix pairs.
15. To show that the inferred coefficients are non-random, we repeated the above steps for sets of shuffled input matrices, using Matlab (R2018a) `randperm` function. We saw that the randomized input matrices result in significantly smaller number of coefficients inferred (Supplementary Fig. 4). We used the `kstest2` function in Matlab to test for the significance in the differences (Kolmogorov-Smirnov distances) between real and shuffled conditions. We obtained p-values = 0 for all comparisons, indicating that the real input has more information content and thus require more coefficients to explain the data.

#### The LOGO module

In addition to the Lasso module, we employ the leave-one-group-out (LOGO) module (Fig. 2d) to obtain a new robust Lasso coefficient matrix of each perturbation in the input dataset. Here, the perturbations are ligand combinations used. We create a ligand-specific matrix by excluding that ligand condition during the model run and comparing the resulting matrix to FULL-data robust Lasso coefficient matrix to determine the coefficients depend on the existence of the corresponding ligand data. We apply LOGO module for both RPPA-RNAseq and RNAseq-ATACseq input pairs. Similar in principle to cross-validation, the LOGO module here enabled us not just to integrate given datasets but to acquire ligand-specific associations.

#### The integration

When both proteomic-transcriptomic and transcriptomic-epigenetic robust Lasso coefficient matrices are obtained, they are merged into a single, gene-level network (Fig. 2c). This network representation is named the Integrated Associations Network (IAN), where each node is a gene, and edge weights represent the Lasso coefficient magnitudes. Notably, the gene nodes can represent data from one or more RPPA, RNAseq, or ATACseq sets. Summarizing the networks at the gene level enabled us to explore pathway enrichments using GSEA (73).

#### GSEA and pathway enrichments

Using the Lasso+LOGO modules and excluding one ligand condition out at a time, we obtain seven LOGO IANs (PBS, EGF, HGF, OSM, IFNγ, TGFβ1, and BMP2) in addition to the FULL-data network. We compare each ligand network with the FULL-data network to determine ligand-dependent associations and create gene-level network visuals using Cytoscape (127). Next, we calculate the weighted sum of edge weights for each gene node in the networks and rank them (Fig. 2e). Then, we run pathway enrichment analysis using GSEA (73) and Reactome to obtain a list of curated pathways enriched for each network (i.e., ligand-LOGO condition).

We calculate the gene-level weight α for each gene k (Equation 2) by summing over each edge width and normalizing by the total number of possible edges.

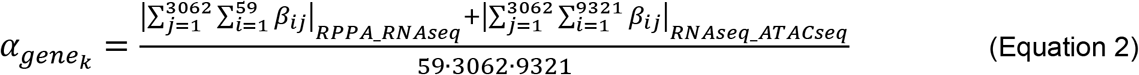

The ranked gene lists are then imported into GSEA software (version 4.1.0) and used in the GSEAPreranked analysis. We select Reactome (v7.2) as the gene set database, keep the default 1000 permutations option, and keep dataset as is without collapsing the gene symbols since we already use the HGNC identifiers. We also choose classical weighting option and choose “149” as our seed for permutation for reproducibility. We repeat these steps for every ligand-specific ranked gene list from the ligand-specific IANs.

After the enrichment analysis successfully completes, we use Enrichment Map Visualization tool of GSEA, together with Cytoscape (v3.7.1). We only retain pathways enriched with a p-value less than 0.05 and false discovery rate (FDR) of 0.1. The results for BMP2 and TGFβ1 are given in Supplementary Data 12 and 13. For this comparison we only retain coefficients with magnitudes larger than 0.1.

### Experimental methods

#### Cell culture

MCF10A cells (ATCC #CRL-10317, acquired from LINCS Consortium/Gordon Mills and STR verified internally in March 2019) are cultured in DMEM/F12 (Gibco #11330032) medium supplemented with 5% (by volume) horse serum (Gibco #16050122), 20ng/mL EGF (PeproTech #AF-100-15), 0.5 mg/mL hydrocortisone (Sigma #H-0888), 10μg/mL insulin (Sigma #I-1882), 100ng/mL cholera toxin (Sigma #C-8052), and 2mM L-Glutamine (Corning #25-005-CI). Cells were cultured at 37^°^C in 5% CO_2_ in a humidified incubator and passaged every 2-3 days with 0.25% trypsin (Corning #25-053-CI) to maintain subconfluency. Experimental starvation medium is DMEM/F12 medium supplemented with 5% (by volume) horse serum (Gibco #16050122), 0.5 mg/mL hydrocortisone (Sigma #H-0888), 100ng/mL cholera toxin (Sigma #C-8052), and 2mM L-Glutamine (Corning #25-005-CI).

#### Mycoplasma testing

The cells were tested for Mycoplasma using a detection kit (Lonza #LT07-701). Following the manufacturer protocol, growth media (2mL) from culture plate was spun at 200g for 5 minutes and 100 μL from the cleared supernatant was transferred into a well of a 96-well plate. 100 μL testing Reagent was added into the same well and let settle for 5 min. Then luminescence is measured in Synergy H1 microplate reader (Agilent Technologies, Inc., CA, USA) with Gen5 (v3.08) software. The Gain was set to 200, Integration time to 1 second, and a single reading was captured. Then, the plate is returned under the hood and 100 μL of testing Substrate was added into the same well. The plate was let settle for 10 min at room temperature. Then, the second luminescence reading (cell supernatant, Reagent, and Substrate) was taken with the same parameters. A ratio of less than 1 indicates a negative test.

#### Validation experiments

The cells were seeded in full growth media at 2000 cells/well in tissue culture treated (no collagen-coating) 96-well plates (Falcon #353072). After loading, plates were left to settle under the hood for 30 minutes and then placed in the incubator for 10 hours. Next, the media is exchanged to experimental starvation media for 15 hours. Then, media are replaced with fresh experimental media containing the ligand(s): EGF (10 ng/mL, R&D Systems #236-EG), BMP2 (20 ng/ml, R&D Systems #355-BM) + EGF (10 ng/ml), and TGFβ1 (10 ng/ml, R&D Systems #240-B) + EGF (10 ng/ml). Each condition was repeated in triplicate. The plates are incubated for ~48 hours. The cells are fixed using %2 paraformaldehyde (Alfa Aesar #43368) and stained with Hoechst (1:10000, by volume, BD Biosciences #561908) for nucleus localization. The plate was left for 1 hour at room temperature. The wells are washed once with 1X PBS and replenished with 40 μl/well PBS.

#### Imaging

The plates are imaged at 10X magnification with phase contrast objective (Agilent/BioTek part number 1320516) and TagBFP filer cube (Agilent/BioTek part number 1225115, excitation 390nm, emission 447nm). A total of 10×8 fields of view per well? are imaged with laser autofocus on the Cytation5 (Agilent Technologies, Inc., CA, USA). When reading was done, the tiles were montaged together by the Gen5 (v3.04, Agilen/BioTeK) software using phase contrast images as registration template (fusion method=linear blend, final image reduction to 13.71%). The imaging parameters for phase contrast were LED=10, Integration time=8, and Gain=24. The parameters for the TagBFP channel were LED=10, Integration time=36, and Gain=24.

#### Image processing and quantification

The images are processed for cell segmentation and finding cell centroids. In short, TrackMate (v7.1.0) plugin from ImageJ (v2.3.0/1.53f) is used to locate cell nuclei in each image (Hoechst stain + BFP channel, see Source Data) and the summary was exported as a comma separated file (see Source Data). The parameters for the plugin were Detector=LoG, Estimated object diameter=5 pixels, Quality threshold=0, Pre-process with median filter=ON, Sub-pixel localization=ON, and Initial thresholding=Auto.

#### Spatial and microenvironmental metric calculations

The csv files exported by ImageJ included each cell object as a row and reported its center coordinates with other default information. The files were imported, and the cells-per-cluster and distance-to-neighbors metrics were evaluated using R scripts (Source Data, github.com/cerdem12/MOBILE), adapted from (github.com/MEP-LINCS/MDD/blob/master/R/MDD_Immunofluorescence_Lvl0Data_Processing.R (67)).

## Code Availability

The final model scripts, files, and information are available at github.com/cerdem12/MOBILE.

## Data Availability

The LINCS datasets analyzed during the current study are available in the Synapse repository, synapse.org/LINCS_MCF10A (67,128). Source Data are provided with this paper at doi:10.6084/m9.figshare.20294229.

## Author Contributions

Conceptualization, C.E. and M.R.B.; Methodology, C.E., S.M.G., L.M.H., and M.R.B.; Software, C.E.; Validation: C.E. and M.R.B.; Formal analysis: C.E.; Investigation: C.E., S.M.G., and L.M.H.; Resources: L.M.H. and M.R.B.; Data curation: C.E. and S.M.G.; Writing – Original Draft: C.E. and M.R.B.; Writing – Review & Editing: C.E. and M.R.B.; Visualization: C.E. and S.M.G.; Supervision: L.M.H. and M.R.B.; Project administration: C.E., L.M.H, and M.R.B.; Funding acquisition: L.M.H. and M.R.B.

## Acknowledgments

The authors acknowledge funding from the National Institutes of Health Grants R01GM104184 (M.R.B.), 1R35GM141891 (M.R.B.), U54HG008098-LINCS Center (M.R.B.), U54CA209988 (L.M.H.), and U54HG008100-LINCS Center (L.M.H.). C.E. was an NIH-LINCS Postdoctoral Fellow.

