## Supplementary material for "Multi-Omics Binary Integration via Lasso Ensembles (MOBILE) for identification of context-specific networks and new regulatory mechanisms": MOBILEintegrator_SuppFigs.pdf

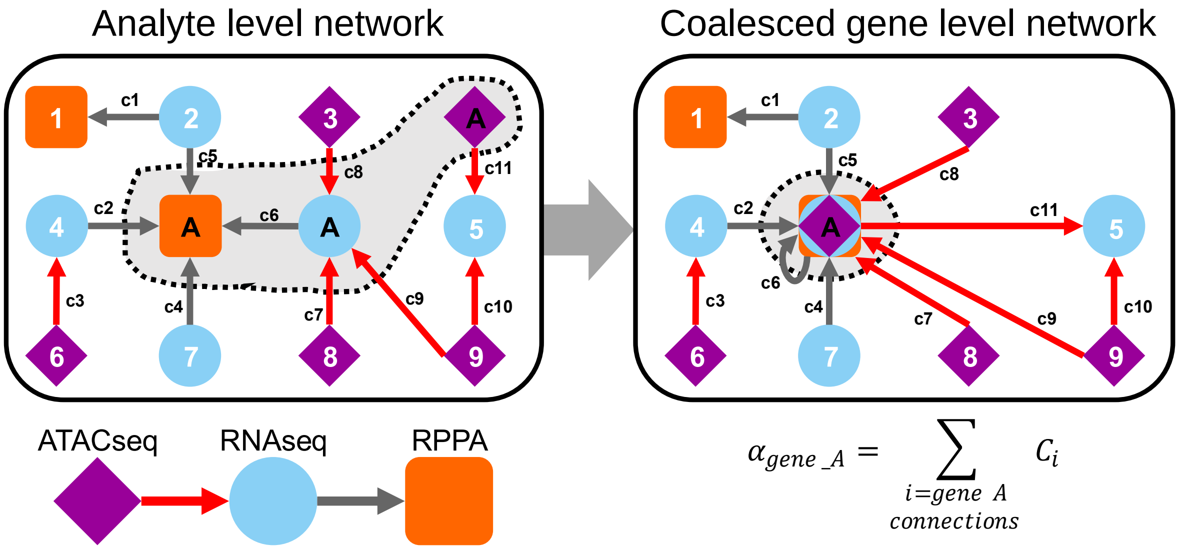


**Supplementary Fig. 1 The analyte level association matrices are used to create gene-level coalescence networks.** Different analytes (ATACseq: diamond, RNAseq: circle, RPPA: square) can have the same gene. For instance, gene A has protein, RNA, and ATACseq peak nodes, each associated with one or more other nodes. The numbers (1-9) represent different nodes corresponding to different genes besides gene A. The analytes with the same gene (i.e., A) are coalesced into a single node, preserving all the edges. After coalescence of all such nodes, the weights (MOBILE inferred association coefficients) of all incoming edges are summed to create a ranked gene-list of the network. Red arrows represent association edges between ATACseq-RNAseq analytes, and the black arrows represent association edges between RNAseq-RPPA nodes. The RPPA analytes can represent phospho- or total protein level measurements.

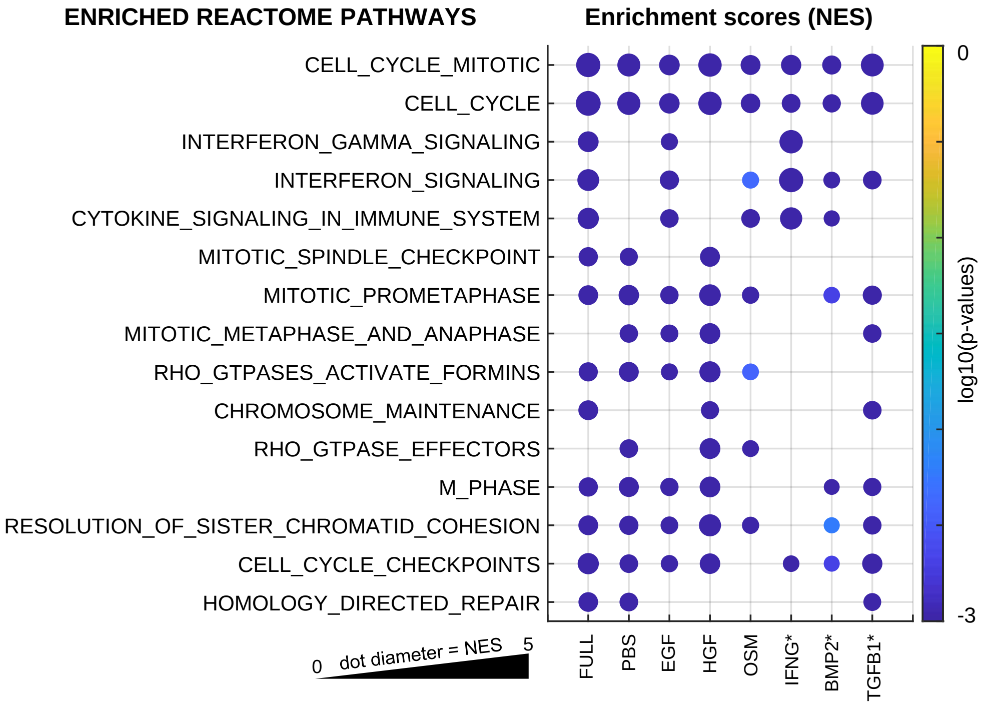


**Supplementary Fig. 2 MOBILE inferred IANs are enriched for canonical pathways and the top associations are literature verified interactions.** Gene set enrichment analysis using genes of FULL and ligand dependent associations reveal cell cycle, interferon, and cytokine signaling pathways. Only top 15 significantly enriched pathways (ranked based on mean scores across conditions) are shown (p-values < 0.05, FDR<0.1). The dot diameter corresponds to the normalized enrichment scores (NES) and color denotes p-values. * denotes conditions with additional EGF treatment.

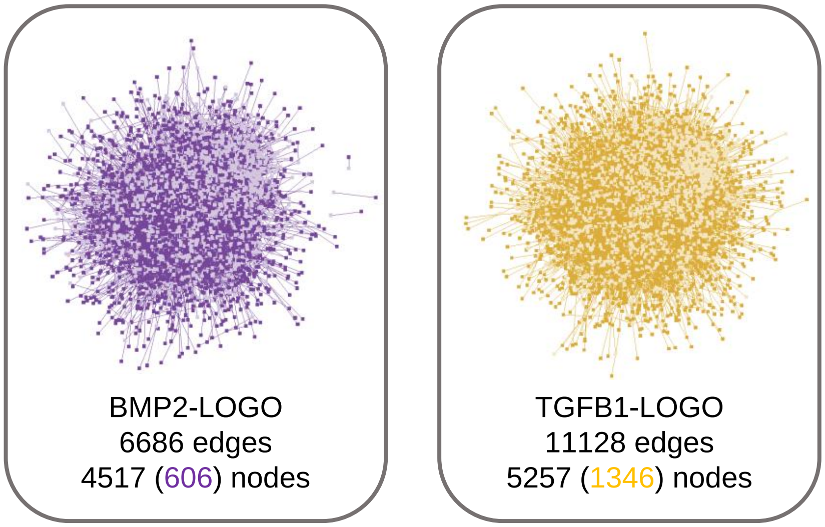


**Supplementary Fig. 3 Comparison of TGFβ1- and BMP2-IANs to FULL network reveals distinct nodes & edges.** There are thousands of ligand-specific associations (number of edges) when the Integrated Association Networks (IANs) of the TGFβ1 and BMP2 are compared. The genes within these sub-networks are ranked by weighted sum of edge magnitudes. Colored numbers represent the number of ligand-specific genes (nodes) in the corresponding networks.

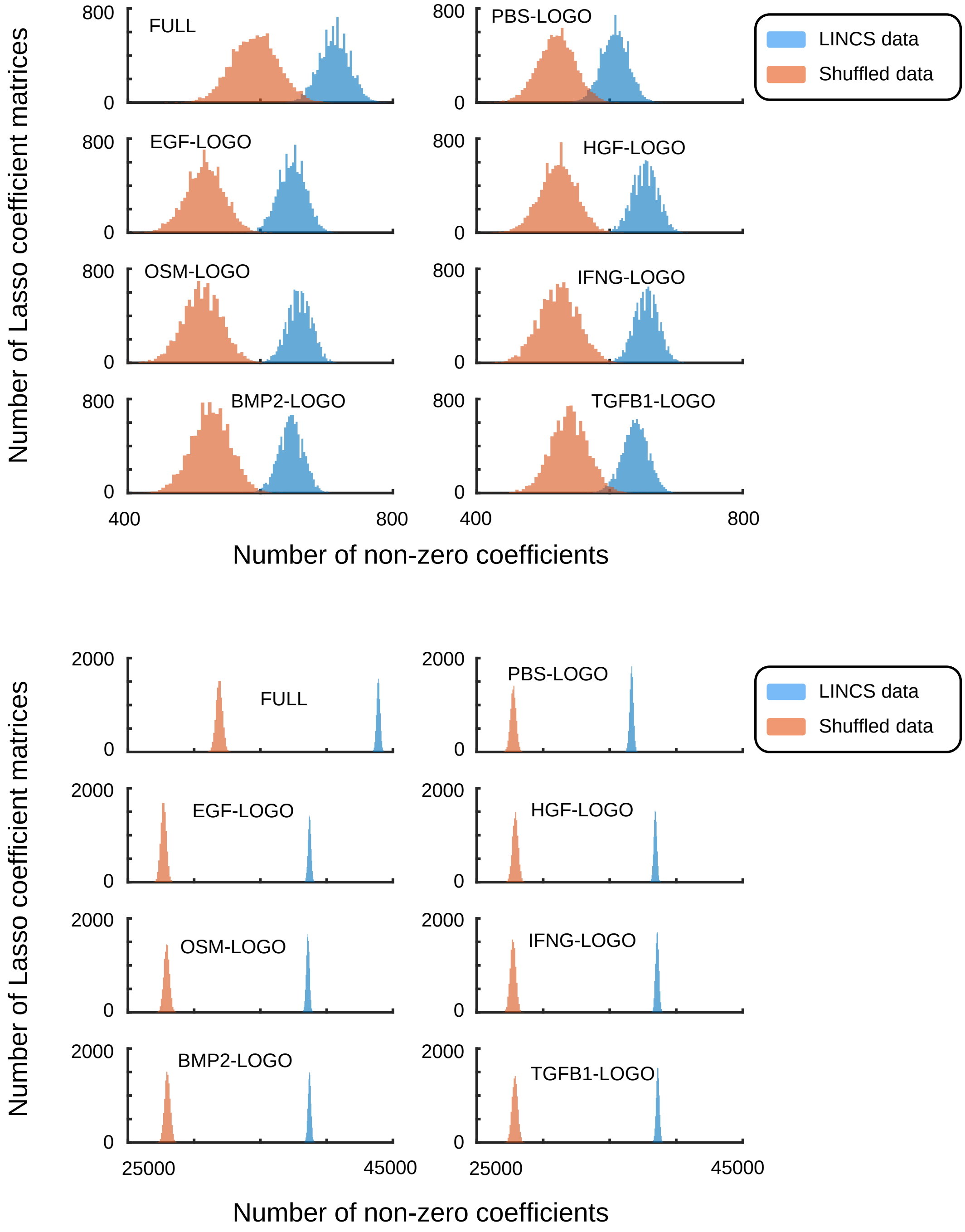


**Supplementary Fig. 4 MOBILE infers significantly different number of Lasso coefficients in real (LINCS) data compared to shuffled (randomized) input. a** Histograms of the Lasso coefficient matrices and their non-zero coefficients from RNAseq-RPPA pair of inputs. Real data input yields more non-zero coefficients inferred, partly due to larger information content. Also note that the FULL data input histograms (top left) are shifted to the right compared to the LOGO conditions. b Histograms from the ATACseq-RNAseq pair of input matrices.
